# Hyaluronic Acid Fuels Pancreatic Cancer Growth

**DOI:** 10.1101/2020.09.14.293803

**Authors:** Peter K. Kim, Christopher J. Halbrook, Samuel A. Kerk, Stephanie Wisner, Daniel M. Kremer, Peter Sajjakulnukit, Sean W. Hou, Galloway Thurston, Abhinav Anand, Liang Yan, Lucia Salamanca-Cardona, Samuel D. Welling, Li Zhang, Matthew R. Pratt, Kayvan R. Keshari, Haoqiang Ying, Costas A. Lyssiotis

## Abstract

Rewired metabolism is a hallmark of pancreatic ductal adenocarcinomas (PDA). Previously, we demonstrated that PDA cells enhance glycosylation precursor biogenesis through the hexosamine biosynthetic pathway (HBP) via activation of the rate limiting enzyme, glutamine-fructose 6-phosphate amidotransferase 1 (GFAT1). Here, we genetically ablated GFAT1 in PDA cell lines, which completely blocked proliferation in vitro and led to cell death. In contrast, GFAT1 knockout did not impair tumor growth, suggesting that cancer cells can maintain fidelity of glycosylation precursor pools by scavenging nutrients from the tumor microenvironment. Here, we show that hyaluronic acid (HA), an abundant carbohydrate polymer in pancreatic tumors composed of repeating N-acetyl-glucosamine (GlcNAc) and glucuronic acid sugars, can bypass GFAT1 to refuel the HBP via the GlcNAc salvage pathway. Furthermore, HA facilitates proliferation in nutrient-starved wild-type PDA. Together, these data show HA can serve as a nutrient fueling PDA metabolism beyond its previously appreciated structural and signaling roles.

## Introduction

Pancreatic ductal adenocarcinoma (PDA) is one of the deadliest human cancers with no clinically effective treatment options (1). PDA is characterized by an intense fibroinflammatory stroma, poor vascularity, low nutrient levels, and rich deposition of extracellular matrix components. To survive and proliferate in this nutrient austere tumor microenvironment, the signature-driving oncogene in PDA, mutant Kras, facilitates the rewiring of PDA metabolism (2-4).

Among the rewired pathways, we previously demonstrated that mutant Kras promotes the activity of the hexosamine biosynthesis pathway (HBP) by upregulating expression of the rate-limiting enzyme glutamine-fructose 6-phosphate amidotransferase 1 (GFAT1) (5). The HBP is an evolutionarily conserved pathway that integrates glucose, glutamine, fatty acid, and nucleotide metabolism to generate the final product uridine diphosphate N-acetylglucosamine (UDP-GlcNAc). UDP-GlcNAc is a crucial donor molecule for glycosylation and O-GlcNAcylation, two essential post-translational modifications required for cellular structure, signaling, and survival (6). The HBP is the only way to generate UDP-GlcNAc *de novo*. Because the HBP integrates nutrients from several major macromolecular classes to produce UDP-GlcNAc, predictably it also acts a nutrient sensing mechanism for available energy within a cell (7). Indeed, numerous studies across cancer subtypes have demonstrated how HBP activity is enhanced to support tumor survival and growth (8-11) and even immune evasion through alteration of extracellular glycosylation content (12).

A compendium of studies during the last decade have revealed that PDA cells fuel their rewired metabolic programs through nutrient scavenging (5, 13-17). Mechanisms include sustained activation of intracellular recycling pathways (e.g. autophagy), the upregulation of nutrient transporter expression (e.g. carbohydrate, lipid, and amino acid transporters), and the activation of extracellular nutrient scavenging pathways (e.g. macropinocytosis). Further, PDA cells also participate in metabolic crosstalk and nutrient acquisition with non-cancerous cells in the tumor microenvironment (TME), such as cancer-associated fibroblasts (CAFs) and tumor-associated macrophages (TAMs) (18-22). A notable example is the observation that PDA cells can directly obtain nutrients from the CAF-derived extracellular matrix (ECM), such as collagen (17). Taken together, elucidating the interaction of PDA cells with different cell populations and ECM components will be instrumental for delineating deregulated PDA metabolism and improving therapeutic strategies.

A major structural component of the TME is hyaluronic acid (HA), a hydrophilic glycosaminoglycan. HA is ubiquitously present in human tissue, especially in skin, connective tissue, and joints, and it is richly abundant in pancreatic tumors (23). HA is primarily deposited by CAFs and, to some extent, by PDA cells (24, 25). HA avidly retains water, which is responsible for both its lubricating properties and, in PDA tumors, the supraphysiological pressure that impairs vascularity and limits drug penetrance (26, 27). An aspect of HA biology that has not previously been studied is its potential role as a nutrient. This is surprising given that HA is a carbohydrate polymer whose monomeric unit is a disaccharide of glucuronic acid and N-acetyl-glucosamine (GlcNAc).

Herein, we set forth to determine the utility of targeting the HBP in PDA. We found that GFAT1 was required for cell survival in vitro. In marked contrast, GFAT1 knockout tumors readily grew in vivo. Based on this observation, we hypothesized that GlcNAc-containing components of the extracellular matrix could bypass the HBP in vivo by way of the GlcNAc salvage pathway. We demonstrate that HA can be metabolized by PDA cells to support growth by refilling the HBP. In sum, our study identifies HA as a novel nutrient source in PDA and contributes to a growing body of data illuminating the important role of the TME in cancer metabolism.

## Results

### Pancreatic cancer cells require *de novo* HBP fidelity in vitro but not in vivo

Previously, we found that mutant Kras transcriptionally activates GFAT1 expression downstream of MAPK signaling in a murine model of PDA to facilitate HBP activity (5). GFAT1 catalyzes the reaction that generates glucosamine 6-phosphate and glutamate from fructose 6-phosphate and glutamine (**Figure 1A**). In another previous study we demonstrated that PDA cells are dependent on glutamine anaplerosis for proliferation (28). Thus, we hypothesized that inhibiting GFAT1 in PDA would have the simultaneous benefit of blocking two major metabolic pathways that support PDA proliferation, thereby providing a considerable therapeutic window.

**Figure 1.**
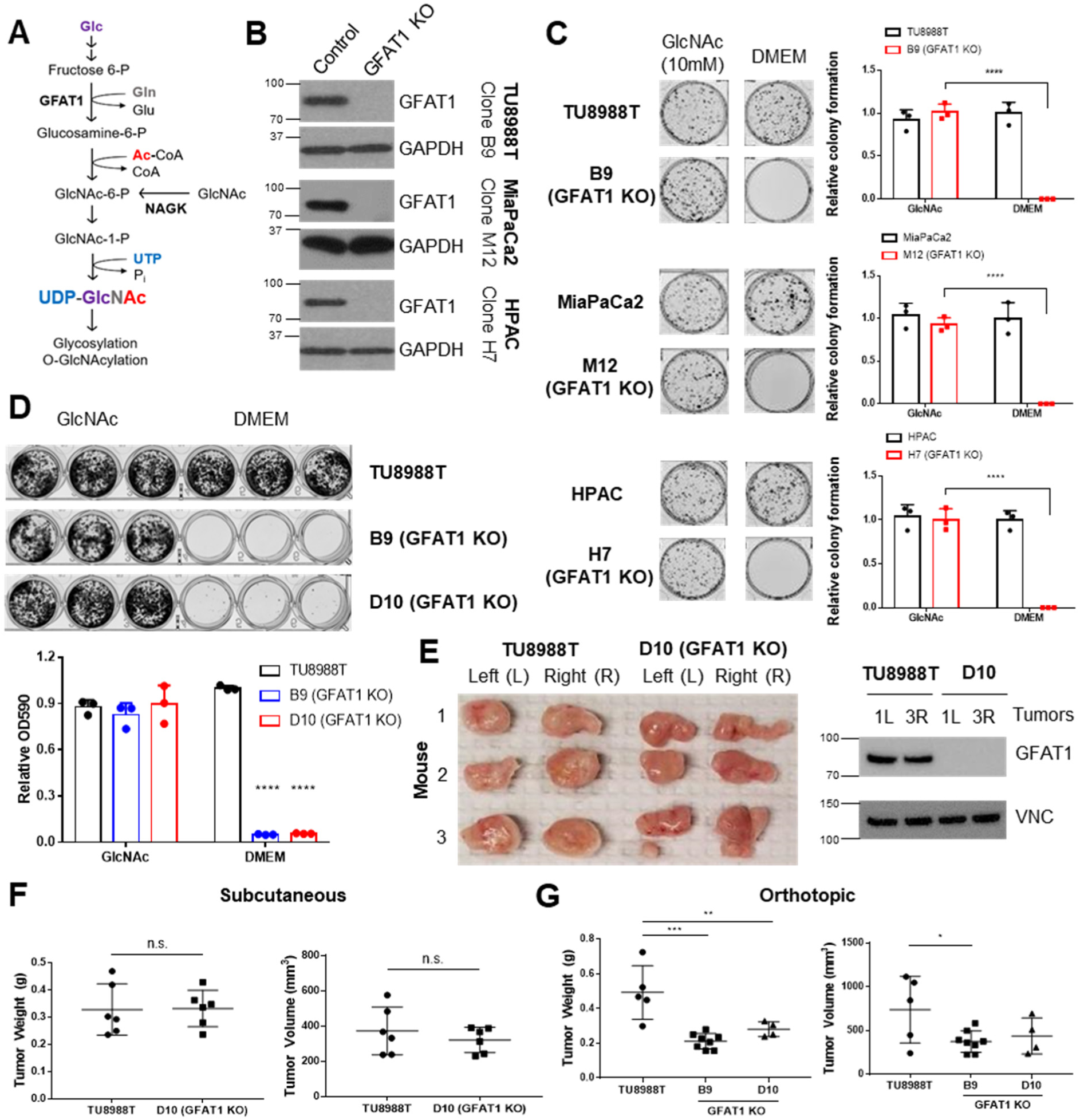
PDA requires de novo hexosamine biosynthetic pathway fidelity in vitro but not in vivo. (**A**) Schematic overview of the hexosamine biosynthetic pathway (HBP) and the nutrient inputs. Ac-CoA, acetyl-coenzyme A; GFAT1, glutamine fructose 6-phosphate amidotransferase 1; Glc, glucose; GlcNAc, N-acetyl-glucosamine; Gln, glutamine; Glu, glutamate; NAGK, N-acetyl-glucosamine kinase; Pi, inorganic phosphorus; UTP, uridine-triphosphate. (**B**) Western blot of GFAT1 and loading control GAPDH from validated CRISPR/Cas9 knockout TU8988T, MiaPaca2, and HPAC clones and their control (non-targeted sgRNA). (**C**) Representative wells from a colony-forming assay in parental and clonally-derived GFAT1 knockout cell lines grown in base media (DMEM) or base media supplemented with 10mM GlcNAc. Data quantitated at right, n=3. (**D**) Proliferation assay in parental and two GFAT1 knockout clonal TU8988T cell lines. Representative wells are presented above data quantitated by crystal violet extraction and measurement of optical density (OD) at 590nm, n=3. (**E**) Tumors from parental TU8988T (n=6) and GFAT1 knockout clone D10 (n=6) grown subcutaneously in immunocompromised mice. Accompanying western blot for GFAT1 and VINCULIN (VNC) loading control from representative tumor lysates. (**F**) Tumor volume and tumor weight from samples in **E**. (**G**) Tumor volume and tumor weight from parental TU8988T (n=5) and GFAT1 knockout clones B9 (n=8) and D10 (n=4) implanted and grown orthotopically in the pancreas of immunocompromised mice. Error bars represent mean ± SD. n.s., non-significant; \**P* < 0.05; ** *P* <0.01; *** *P* <0.001; **** *P* <0.0001.

Our previous results targeting GFAT1 in murine cells with shRNA yielded insufficient knockdown to draw a conclusion as to its necessity in PDA (5). Thus, here we used CRISPR/Cas9 to knock GFAT1 out from three established human PDA cell lines: HPAC, TU8988T, and MiaPaCa2. During selection, the pooled polyclonal populations were grown in GlcNAc, which bypasses GFAT1 via the GlcNAc salvage pathway (**Figure 1A**). This supplement was included to minimize metabolic rewiring within the selected populations.

The GFAT1 knockout lines had differential response to GlcNAc withdrawal. Among the three GFAT1 knockout cell lines, only the HPAC line exhibited a marked reduction in cell number, consistent with loss of viability, in the 4 days following GlcNAc withdrawal (**Supplementary Figure 1A**). The impact on proliferation was consistent with the decrease in the UDP-GlcNAc pool, which was analyzed using liquid chromatography-coupled tandem mass spectrometry (LC-MS/MS) (**Supplementary Figure 1B**). Consistent with the proliferative phenotypes across lines, the HPAC line also had a significantly smaller UDP-GlcNAc pool than that of either MiaPaCa2 or 8988T cells (**Supplementary Figure 1C**). Cellular O-GlcNAc-ylation of the proteome was also measured by immunoblot three days after GlcNAc withdrawal. Again, consistent with the LC-MS/MS analysis, O-GlcNAc expression was significantly reduced in HPAC but was maintained in 8988T (**Supplementary Fig 1D**).

The data from TU8988T and MiaPaca2 were similar to those from our earlier studies (5), and thus we posited that knockout of GFAT1 was incomplete. As such, we subsequently generated clonal cell lines from the pooled lines. This analysis revealed that the degree of GFAT1 knockout varied by cell line and by clone, and this correlated with their differential growth and sensitivity to GlcNAc withdrawal *in vitro* (**Supplementary Figure 1E,F**). Clones for each cell line without detectable GFAT1 expression (**Figure 1B**) were further validated by sequencing and were subsequently used to examine the role of the HBP more accurately.

Using our genomically-sequenced and bona fide GFAT1 knockout clonal lines, we found that GFAT1 knockout led to an abolishment of colony formation (**Figure 1C**) and potently impaired proliferation (**Figure 1D, Supplementary Figure 1G**) in all three PDA cell lines in vitro. We then moved these cells into *in vivo* tumor models. Surprisingly, when either the pooled or the clonal knockout lines were implanted into the flanks of immunocompromised mice, they readily formed tumors that were comparable to their wild type counterparts in terms of weight and volume (**Figure 1E,F** and **Supplementary Figure 1H**). Similar results were obtained for GFAT1 knockout clonal lines implanted orthotopically into the pancreas (**Figure 1G**). Of note, while clearly capable of forming tumors, the GFAT1 knockout clonal lines grown in the pancreas were smaller than the wild type tumors at endpoint. The marked discrepancy in phenotype between *in vitro* and *in vivo* settings led us to hypothesize that GFAT1 knockout clones were scavenging nutrients from the TME to refill the HBP, which enabled their survival and tumor growth.

### Conditioned media rescues proliferation of GFAT1 knockout PDA cells

To test our scavenging hypothesis, we generated conditioned media (CM) from CAFs, the most abundant stromal cell type in the pancreatic TME (29, 30). When GFAT1 KO clones were incubated in patient-derived CAF CM, we observed a significant, albeit modest, rescue in colony formation (**Figure 2A,B**). Unexpectedly, we observed a more robust, dose-dependent rescue of colony formation in GFAT1 knockout cells with CM from wild type TU8988T cells (**Figure 2C-F** and **Supplementary Figure 2A**). Similarly, CM from wild type HPAC and MiaPaCa2 cells was also able to partially rescue proliferation of a subset of GFAT1 KO clones (**Figure 2G** and **Supplementary Figure 2B,C**).

**Figure 2.**
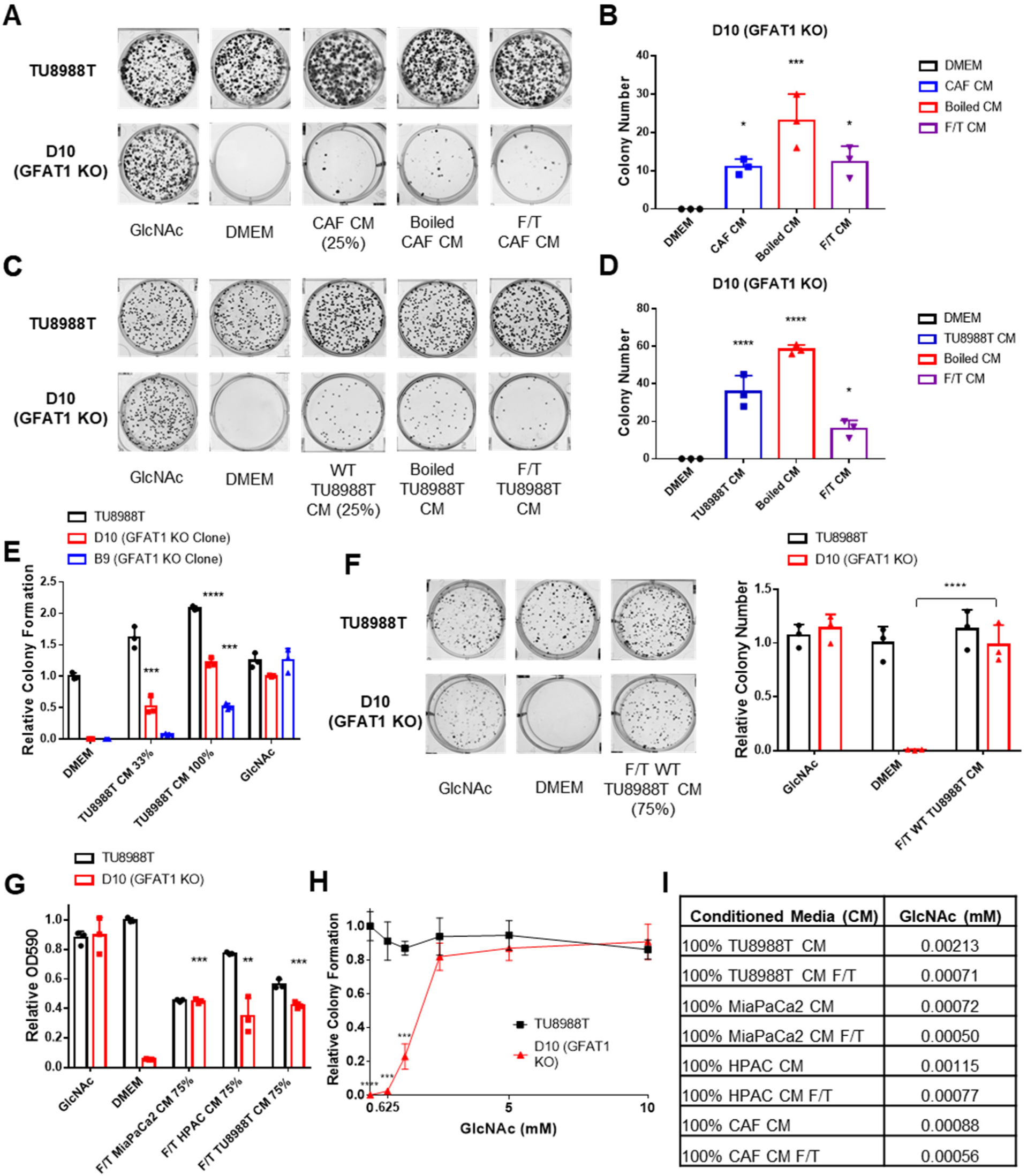
Conditioned media from wild type PDA cells support proliferation of GFAT1 knockout cells. (**A**) Representative wells from a colony-forming assay in parental TU8988T and GFAT1 knockout clonal line D10 in 10mM GlcNAc, base media (DMEM), or base media supplemented 3:1 with cancer associated fibroblast (CAF) conditioned media (CM), boiled CAF CM, or CAF CM subject to freeze-thaw (F/T). (**B**) Quantitation of colonies from data in **A** (n=3). (**C**) Representative wells from a colony-forming assay in parental TU8988T and GFAT1 knockout clonal line D10 in 10mM GlcNAc, DMEM, or base media supplemented 3:1 with CM from wild type TU8988T cells, boiled TU8988T CM, or TU8988T CM subject to F/T. (**D**) Quantitation of colonies from data in **C** (n=3). (**E**) Quantitation of colony forming assay data of parental and GFAT1 knockout clonal TU8988T lines in base media, positive control GlcNAc, wild type TU8988T CM diluted 1:2 (33%) or used directly (100%) (n=3). (**F**) Representative wells and quantitation of colony forming assay data of parental and GFAT1 knockout clonal TU8988T lines in base media, positive control GlcNAc, and wild type TU8988T CM subject to F/T and diluted 3:1 (75%) (n=3). (**G**) Quantitation of colony forming assay data of parental and GFAT1 knockout clonal TU8988T lines in base media, positive control GlcNAc, or wild type TU8988T, HPAC, or MiaPaCa2 CM subject to F/T and diluted 3:1 (75%) (n=3). (**H**) GlcNAc dose response curve presented as relative colony number for parental and GFAT1 knockout TU8988T cells (n=3). (**I**) Absolute quantitation of GlcNAc in various CM by LC-MS/MS (n=3). Error bars represent mean ± SD. n.s., non-significant; *P < 0.05; ** P <0.01; *** P <0.001; **** P <0.0001.

To begin to identify the rescue factors in the CM, we subjected the CM to boiling or repeated cycles of freezing and thawing (F/T). In each of these conditions, both the CAF and the PDA CM retained the ability to support colony formation in GFAT1 knockout cells (**Figure 2A,B**). These results suggested the relevant factor(s) did not require tertiary structure. Additionally, we observed that the rescue activity of the CM was dose dependent (**Figure 2E-G** and **Supplementary Figure 2A-C**).

As GlcNAc was used to establish our GFAT1 knockout lines, we first quantitated the GlcNAc concentration in the CM. GlcNAc dose response curves demonstrated that millimolar quantities of GlcNAc (>0.625mM) were required to rescue colony formation of GFAT1 knockout PDA cells (**Figure 2H** and **Supplementary Figure 2D**). By contrast, LC-MS/MS quantification of GlcNAc in the CM revealed that it was in the low micromolar range (**Figure 2I**), several orders of magnitude below the millimolar doses of exogenous GlcNAc required to maintain proliferation (**Figure 2H** and **Supplementary Figure 2D**). These results illustrated that free GlcNAc was not the relevant molecule in the CM mediating rescue. This led us to consider alternate possibilities, including GlcNAc-containing components of the TME.

### Hyaluronic acid facilitates proliferation in GFAT1 knockout PDA cells and nutrient-starved wild type PDA cells

GlcNAc is a widely utilized molecule as a structural component of the extracellular matrix, a modification of various lipid species, and a post-translational modification on proteins (31, 32). Thus, we hypothesized that GlcNAc was released into CM as a component part of a lipid, protein, or glycosaminoglycan polymer, and that this mediated rescue of GFAT1 knockout. To test this, we first applied necrotic cellular debris from FL5.12 cells (33), which contains the full complement of biomolecules from dead cells, including GlcNAc-containing proteins and lipids, to GFAT1 knockout cells grown at clonal density. Necrotic cell debris was unable to rescue GFAT1 knockout across our cell line panel (**Supplementary Figure 3A-F**). Next, we tested if glycosaminoglycan carbohydrate polymers could mediate rescue of GFAT1 knockout, in a matter akin to CM. High dose heparin was not able to rescue colony formation in GFAT1 knockout cells (**Supplementary Figure 3A-F**), but 78 kDa HA provided a modest but significant rescue (**Figure 3A,B**).

**Figure 3.**
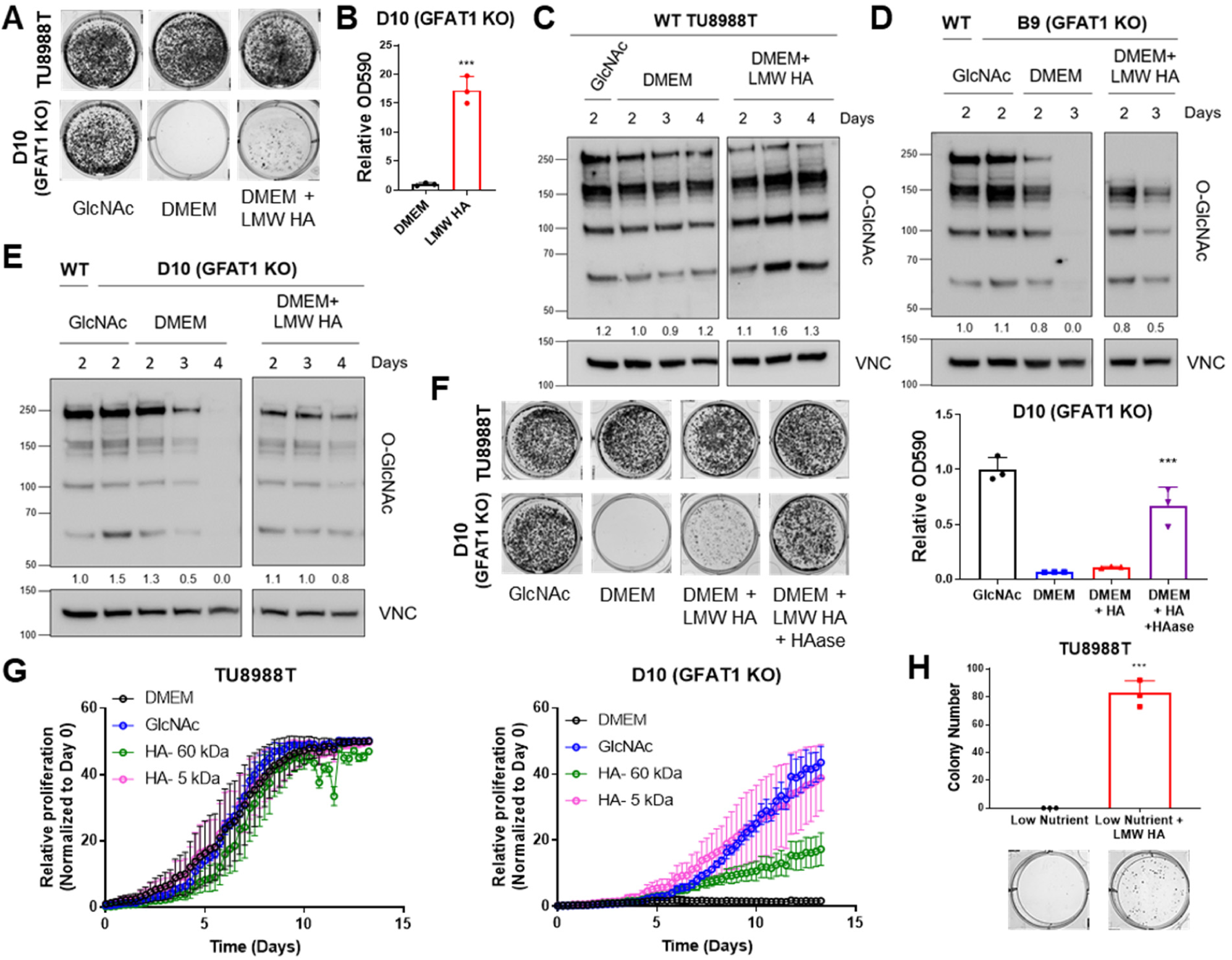
Hyaluronic acid rescues proliferation in GFAT1 knockout PDA cells and nutrient starved PDA cells. (**A**) Representative wells from a colony-forming assay in parental and clonally-derived GFAT1 knockout TU8988T cell lines grown in base media (DMEM), positive control GlcNAc (10mM), or low molecular weight (LMW) hyaluronic acid (78kDa HA, 10mM). (**B**) Quantitation of data from **A** (n=3). (**C**) Western blot of proteome O-GlcNAc and loading control VINCULIN (VNC) in parental TU8988T cells grown in base media (DMEM) plus GlcNAc or LMW HA for the indicated time points. Band density was quantitated, normalized to control, and plotted below the blot. (**D,E**) Western blot of proteome O-GlcNAc and loading control VNC in GFAT1 knockout clonal lines (**D**) B9 and (**E**) D10 in base media (DMEM) plus GlcNAc or LMW HA for the indicated time points. Wild type (WT) TU8988T included as control. Band density was quantitated, normalized to control, and plotted below the blot. (**F**) Representative wells of a proliferation assay in parental TU8988T and GFAT1 knockout clonal line D10 grown in base media (DMEM), positive control GlcNAc (10mM), orbase media supplemented 1:1 with boiled low molecular weight (LMW) hyaluronic acid (HA, 10mM) with and without pre-digestion with hyaluronidase (HAase). At endpoint, cells are stained with crystal violet, and the stain is then extracted and quantitated by OD at 590nm (n=3). (**G**) Proliferation time course, as measured on the Incucyte, of parental TU8988T and GFAT1 knockout cells in base media (DMEM), positive control (GlcNAc), 60 kDa HA (LMW HA), or 5 kDa HA (o-HA) (n=3). (**H**) Quantitated colony forming assay data and representative wells of parental TU8988T cells grown in low nutrient conditions (20-fold reduction in glucose, glutamine, and 10-fold reduction in serum) in the presence or absence of 10 mM LMW HA (n=3). Error bars represent mean ± SD. *** *P* <0.001.

HA is a carbohydrate polymer and an extracellular matrix component that is abundant in the PDA tumor microenvironment (23). The monomeric form of HA is a repeating disaccharide consisting of glucuronic acid and GlcNAc. HA polymer length, often described by its molecular weight (MW), has important impacts on its biological activity. In non-pathological settings, newly synthesized HA is predominantly high molecular weight (HMW; >1000kDa) (34). However, in tumors and tumor interstitial fluid, there is a significantly elevated level of low molecular weight (LMW; 10-250kDa) and oligo-HA (o-HA; <10kDa) (35, 36). Consistent with the rescue of colony formation in GFAT1 knockout cells, LMW HA (78 kDa) was also able to rescue total cellular O-GlcNAc levels, as assessed by western blot (**Figure 3C-E**).

Cancer cells have been reported to uptake HA via macropinocytosis (37). Thus, a possible explanation for the modest rescue could be low macropinocytosis activity. However in PDA, mutant Kras drives high macropinocytosis (13), and quantitation of macropinocytotic activity with a fluorescent dextran-based assay revealed that our three PDA cell lines exhibited considerable macropinocytosis (**Supplementary Figure 3G**). This led us to hypothesize that HA entry into cells is not the rate limiting step, but rather the cleavage of HA into smaller fragments. Consistent with this, breaking down LMW HA with hyaluronidase enhanced the rescue of colony formation (**Figure 3F**). Of note, hyaluronidase was heat-inactivated before its application to GFAT1 knockout cells (**Supplementary Figure 3H**), as hyaluronidase has been reported to directly impact cellular metabolism (38). Next, we tracked the rescue of proliferation by LMW HA (60kDa) and o-HA (5kDa). This analysis revealed that HA-mediated rescue in proliferation was considerably higher for o-HA than for LMW HA (**Figure 3G**).

The studies detailed above were performed with GlcNAc auxotrophs. To determine the effect of HA in a more physiologically relevant setting, we provided HA to wild type PDA cells cultured in low-nutrient media (i.e. a 20-fold reduction in glucose and glutamine, and a 10-fold reduction in serum concentration). Here we found that LMW HA rescued colony formation (**Figure 3H**). Collectively, these data point to a novel role of HA to restore the HBP, promote survival and proliferation of GFAT1 null PDA and, moreover, also that of nutrient-starved wild type PDA cells.

### Hyaluronic acid rescues GFAT1 null PDA via the GlcNAc salvage pathway

The GlcNAc salvage pathway bypasses GFAT1 by catalyzing the phosphorylation of GlcNAc to GlcNAc-6-phsophate, in a reaction mediated by N-acetyl-glucosamine kinase (NAGK). This GlcNAc-6-phosphate is subsequently converted into UDP-GlcNAc (**Figure 1A**). Therefore, we hypothesized that the carbohydrate polymer HA, which is 50% GlcNAc, fuels the HBP via the GlcNAc salvage pathway through NAGK. To test this, we employed the same CRISPR/Cas9 strategy to target NAGK (**Figure 4A**). Knockout of NAGK in parental TU8988T and MiaPaCa2 cell lines had no impact on colony formation, while reducing the colony forming capacity of HPAC cells (**Figure 4B,C**). These results were consistent with the elevated expression of NAGK in HPAC cells (**Figure 4D**). Of note, NAGK knockout did not result in up-regulation of GFAT1 (**Figure 4A**), which could have suggested a compensatory metabolic rewiring.

**Figure 4.**
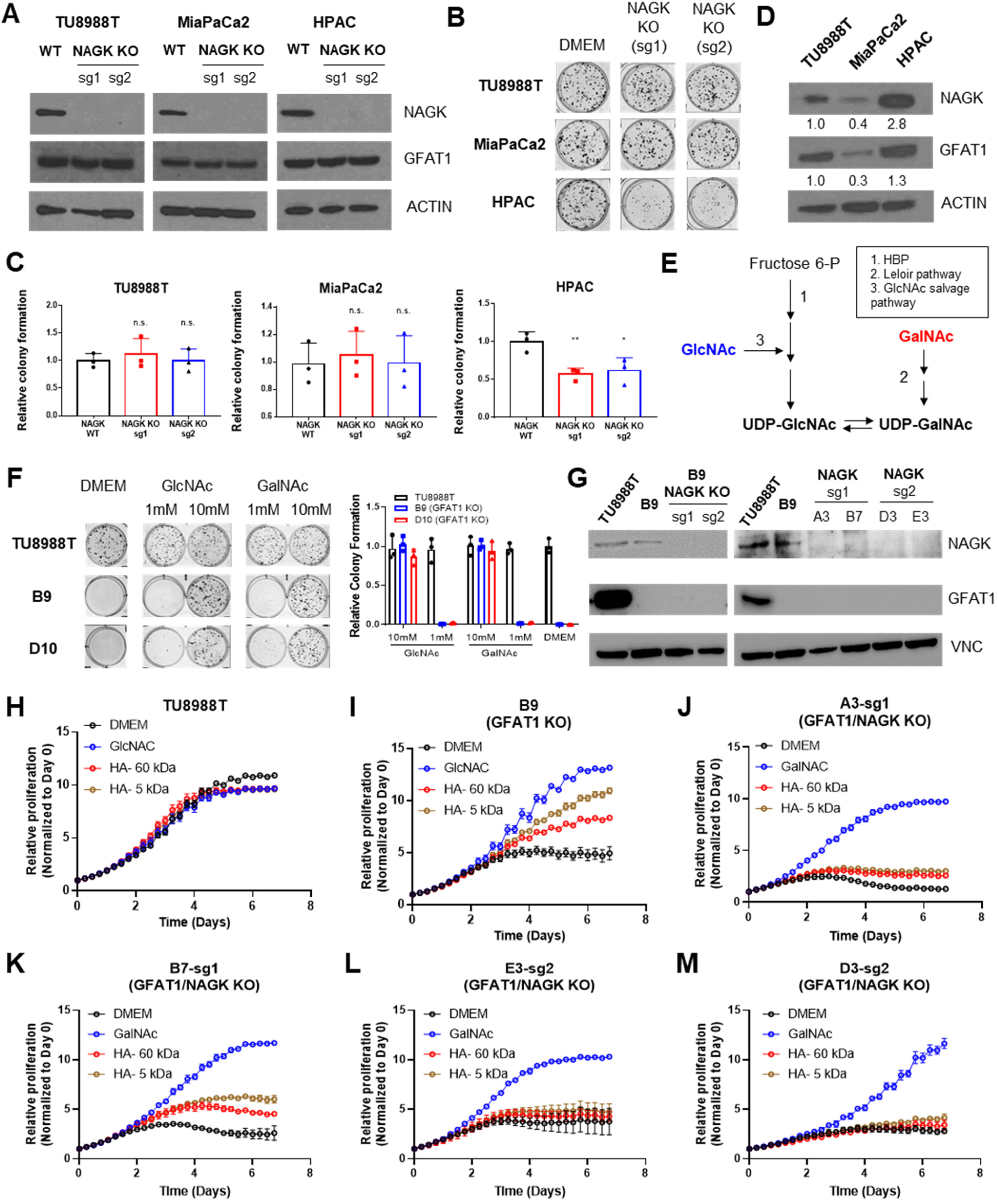
Hyaluronic acid-derived GlcNAc rescues GFAT1 loss via the GlcNAc salvage pathway. (**A**) Western blot of NAGK, GFAT1, and ACTIN loading control from TU8988T, MiaPaCa2, and HPAC parental (wild type, WT) and NAGK KO populations. NAGK was knocked out using two independent sgRNAs (sg1, sg2). (**B**) Representative wells from a colony forming assay for parental and NAGK knockout lines. (**C**) Quantitation of colony forming assay data in **B** (n=3). (**D**) Western blot for NAGK, GFAT1, and loading control ACTIN in parental PDA cell lines. Band density was quantitated, normalized to control, and plotted below the blot. (**E**) Schematic overview of the Leloir pathway of galactose catabolism integrated with the HBP and GlcNAc salvage pathway. (**F**) Representative wells from colony formation assays in parental and GFAT1 knockout clonal TU8988T cell lines in base media (DMEM), positive control GlcNAc, and N-acetyl-galactosamine (GalNAc). (**G**) Western blot for GFAT1, NAGK, and loading control VINCULIN (VNC) in parental TU8988T and HPAC, GFAT1 knockout clones, and GFAT/NAGK double targeted lines. (**H-M**) Proliferation time course, as measured on the Incucyte, of parental (**H,I**) TU8988T and GFAT1 knockout line B9 in base media, GlcNAc positive control, 60 kDa HA, or 5 kDa HA; (**J-M**) GFAT1/NAGK double targeted clones in base media, GalNAc positive control, 60 kDa HA, or 5 kDa HA (n=3). Error bars represent mean ± SD. n.s., non-significant; \**P* < 0.05; ** *P* <0.01.

Next, we targeted NAGK in our GFAT1 knockout clones. GFAT1/NAGK double knockout cells were generated in media containing N-acetyl-galactosamine (GalNAc), an isomer of GlcNAc. Supplementation with GalNAc enables bypass of both the *de novo* HBP and the GlcNAc salvage pathway, by way of the Leloir pathway (39), to support UDP-GalNAc and ultimately UDP-GlcNAc biogenesis (**Figure 4E**). In this way, we were again able to select viable lines while avoiding the selection of those with unpredictable metabolic adaptations.

The GalNAc dose response for GFAT1 knockout clones was comparable to that of GlcNAc (**Figure 2H**), demonstrating that they are indeed viable in GalNAc (**Figure 4F**). Although NAGK expression was efficiently knocked down in our pooled populations (**Figure 4G**), we again selected for clones in order to minimize the effect of NAGK-proficient clones persisting in the bulk population. From among these, we selected four GFAT1/NAGK clones and tracked their proliferation using the Incucyte upon rescue with varying sizes of HA or GalNAc (**Figure 4H-M**). These were compared relative to wild type TU8988T cells (**Figure 4H**) and the GFAT1 knockout line (**Figure 4I**). In stark contrast to the GFAT1 knockout line, LMW HA-and o-HA was unable to rescue GFAT1/NAGK double knockout lines (**Figure 4I-M**). These results illustrate that HA rescue requires NAGK and the GlcNAc salvage pathway, consistent with the idea that HA-derived GlcNAc fuels UDP-GlcNAc biosynthesis upon GFAT1 knockout. Altogether, our data implicate HA as a novel nutrient for PDA. Mechanistically, HA regulates PDA metabolism by refueling the HBP via the GlcNAc salvage pathway, supporting PDA survival and proliferation.

## Discussion

The HBP is activated in a Kras-dependent manner in PDA (5), and it is similarly elevated in numerous cancers to provide a diverse set of functions, including the regulation of proliferation, survival, angiogenesis, and metastasis (9). As such, we and others have proposed that the HBP may provide a selective vulnerability for cancer therapy, with GFAT1 as an attractive therapeutic target (5, 10, 40, 41). Indeed, several pan glutamine-deamidase inhibitors (e.g. azaserine and 6-diazo-5-oxo-L-norleucine), which potently suppress GFAT activity, have demonstrated anti-tumor activity *in vitro* and *in vivo* in PDA and other cancers (42, 43). However, because these drugs are not specific to the HBP, it has not been clear what impact GFAT-specific inhibition had on these phenotypes. As such, we took a genetic approach to knock out GFAT1 to elucidate the role of the HBP in PDA.

In the PDA models tested herein, GFAT1 knockout was not compatible with PDA cell proliferation *in vitro*, unless the media were supplemented with GlcNAc or GalNAc (**Figure 1C,D** and **Figure 4F**). However, these same cells readily formed tumors *in vivo* in subcutaneous and orthotopic models (**Figure 1E-G**). The stark discrepancy in phenotypes led us to hypothesize that the TME was providing the means to bypass GFAT1. Indeed, we found that denatured conditioned media from CAFs and wild type PDA cells were able to rescue viability in GFAT1 knockout PDA cells, implicating a molecule(s) without tertiary structure (**Figure 2**). By examining several GlcNAc-containing candidates, we discovered a previously unknown role of HA as a nutrient source for PDA (**Figure 3**). We report that HA can refill the HBP via the GlcNAc salvage pathway (**Figure 4**) to support PDA survival and proliferation.

HA is traditionally regarded as a structural component in physiology (44). In addition to this role, a wealth of studies have ascribed additional functions to HA. For example, HA can activate cell-cell contact-mediated signal transduction through CD44 and/or receptor for HA-mediated motility (RHAMM) (45). The signaling activity/function of HA depends on its MW (44, 46). Similarly, a recent study illustrated that breakdown of the HA matrix with hyaluronidase enabled the interaction between growth factors and growth factor receptors (38). This promoted glucose metabolism, cellular proliferation, and migration. The role of HA in GFAT1 knockout and nutrient starved PDA cells described herein is likely independent of its structural and signaling functions, given that we observe considerably greater rescue with o-HA (**Figure 3F,G**), a form of HA that is not traditionally considered for these purposes.

Rather, our study introduces a novel role to HA as a fuel for PDA tumor growth (**Figure 3G,H**), further highlighting the significance and biological complexity of this predominant glycosaminoglycan. Additionally, our study suggests that NAGK, through which HA-mediated GlcNAc presumably refuels the HBP in vivo, may be an attractive therapeutic target for PDA. Indeed, a recent study demonstrated that NAGK knockout in PDA impairs tumor growth in vivo, while only exhibiting a modest impact on cellular proliferation in vitro (*Wellen lab, co-submitted study*). These results are consistent with our observations on utilization of the GlcNAc salvage pathway to fuel UDP-GlcNAc pools from HA-derived GlcNAc (**Figure 4E**). Our study also contributes to a growing body of data illuminating unexpected nutrient sources in the TME that support cancer metabolism (13, 14, 16-21, 47), and this raises the possibility that other glycosaminoglycans may be similarly scavenged.

Due to its extremely hydrophilic nature, HA retains water and acts as a cushioning agent in tissue homeostasis and biomechanical integrity (44). In PDA, HA is a predominant component of the TME, and its water-retaining property is one of the main drivers of the supraphysiological intratumoral pressure (48). This pressure can exceed 10-fold that observed in the normal pancreas, and, as a result, tumor vasculature collapses (49-51). The limited access to circulation impairs nutrient and oxygen delivery, and it has been proposed that this is a critical impediment to tumoral drug delivery (52). Indeed, in animal models, breakdown of the HA matrix with a therapeutic hyaluronidase (PEGPH20) reduces intratumoral pressure, restores circulation, which facilitates drug delivery, and thereby improves response to chemotherapy (50, 51). Based on these promising observations, PEGPH20 was tested in clinical trials alongside standard of care chemotherapy. Despite the successes in the preclinical models, PEGPH20 did not extend PDA patient survival (53).

The discrepancy between the clinical response to PEGPH20 and the preclinical data remains an active area of investigation and may concern the myriad additional roles of HA. For example, the HA matrix may be necessary to restrain tumor dissemination, as was shown for CAF depletion studies in PDA (54-57). Thus, the benefits afforded by enhanced drug penetration facilitated by PEGPH20 may be negated by this side effect. Along these lines, HA degradation may also enhance tumor metabolism and growth. This could occur through growth factor signaling-dependent (38) as well as signaling-independent pathways, like the GlcNAc salvage pathway described herein. In contrast, reduction in the HA content of tumors also facilitates T cell invasion (43), which may complement immunotherapy approaches, a concept that would be hindered by immunosuppressive chemotherapies. Given the conflicting roles of HA in tumor restraint and tumor growth, considerable work remains to be done to determine the most effective way to exploit this feature of pancreatic cancer.

## Materials and Methods

### Cell Culture

MiaPaCa2 and HPAC were obtained from ATCC. TU8988T was obtained from DSMZ. Patient-derived CAFs (58) were a generous gift from Rosa Hwang, and FL5.12 cells were a generous gift from Dr. Aimee Edinger. All cells were routinely checked for mycoplasma contamination with MycoAlert PLUS (Lonza) and validated for authenticity annually by STR profiling. Cells were maintained in standard high glucose DMEM without pyruvate (Gibco) supplemented with 10% fetal bovine serum (FBS; Corning). GFAT1 null PDA were cultured in standard media supplemented with 10mM GlcNAc (Sigma). GFAT1 null NAGK knockout PDA were cultured in standard media supplemented with 10mM GalNAc (Sigma). Low nutrient media was made with DMEM without glucose, glutamine and pyruvate (Gibco). Glucose, glutamine, and FBS were added to the final concentration of 1.25mM, 0.2mM and 1%, respectively. FL5.12 cells were maintained in RPMI 1640 (Gibco) supplemented with 10% FBS, 10mM HEPES (Sigma), 55μM β-mercaptoethanol (Sigma), antibiotics, 2mM glutamine, and 500 pg/mL recombinant murine IL-3 (Peprotech 213-13).

### Generation of CRISPR/Cas9 knockout clones

GFAT1 and NAGK knockout PDA cell lines were generated using CRISPR/Cas9 method described previously (22). Overlapping oligonucleotides (Feng Zhang lab human GeCKOv2 CRISPR knockout pooled library; Addgene #1000000048) were annealed to generate sgRNA targeting GFAT1 or NAGK. sgRNA was cloned directly into the overhangs of PX459 V2.0 vector (Feng Zhang lab; Addgene plasmid #62988) that was digested with BbsI. The resulting CRISPR/Cas9 plasmid was transformed into chemically competent Stbl3 cells, miniprepped for plasmid DNA, and sequence-verified. sgRNA oligonucleotide pairs for GFAT1 (10) and NAGK are as follows: GFAT1 (sg1 Fwd 5′-CACCgCTTCAGAGACTGGAGTACAG-3′; sg1 Rev 5′-AAACCTGTACTCCAGTCTCTGAAGc-3′) and NAGK (sg1 Fwd 5’-CACCgTAGGGGAGGCACACGATCCG; sg1 Rev 5’-AAACCGGATCGTGTGCCTCCCCTAc-3’; sg2 Fwd 5’-CACCgGCCTAGGGCCTATCTCTGAG-3’; sg2 Rev 5’-AAACCTCAGAGATAGGCCCTAGGCc-3’). Human PDA were transiently transfected using Lipofectamine 3000 according to the manufacturer’s instructions. Cells were selected with puromycin in the presence of GlcNAc (GFAT1 knockout bulk population) or GalNAc (GFAT1 NAGK double knockout bulk population). To select clones, polyclonal pools were seeded into 96-well plates at a density of 1 cell per well. Individual clones were expanded and verified via western blot and Sanger sequencing. TU8988T clone B9 has a 10 base pair (bp) and a 1bp deletion in GFAT1; TU8988T clone D10 has 2 different 1bp deletions in GFAT1; MiaPaCa2 clone M12 has 2 different 1bp deletions in GFAT1; HPAC clone H1 has a 187bp deletion in GFAT1; HPAC clone H7 has a 187bp deletion in GFAT1.

### Conditioned media

Conditioned media was generated by culturing cells in 15 cm^2^ plates (25mL growth media/plate) for 72 hours at 37°C, 5% CO_2_, so that they reached ∼90% confluence. The media were then filtered through a 0.45μm polyethersulfone membrane (VWR). Boiled conditioned media was warmed to 100°C for 15 minutes. Freeze-thaw conditioned media were initially stored at −80°C and were thawed in a 37°C water bath on the day of the experiment. As indicated, fresh growth media were added to the conditioned media at the ratios indicated to avoid nutrient/metabolite exhaustion.

### Colony formation and proliferation assays

For colony formation assays, cells were plated in a 6-well plate in biological triplicate at 500 cells/well in 2 mL of media and grown for 9-12 days. For proliferation assays, 5000 cells/well were plated. At end point, assays were fixed with 100% methanol for 10 minutes and stained with 0.5% crystal violet solution for 15 minutes. Relative colony formation was quantitated manually in a blinded fashion. Proliferation was quantified by removing the dye with 10% acetic acid and measuring OD595.

### CyQUANT viability assay

Cells were seeded in 96-well black wall, clear bottom plates at 1000 cells/well in 50μL of media and incubated at 37°C, 5% CO_2_ for indicated time points. At each time point, media was aspirated and plates were stored at −80°C. Proliferation was determined by CyQUANT (Invitrogen) according to the manufacturer’s instructions. SpectraMax M3 Microplate reader (Molecular Devices) was used to measure fluorescence.

### IncuCyte S3: Real-time, live-cell proliferation assay

1000 cells were seeded per well in a 96-well plate and incubated at 37°C, 5% CO_2_ for cells to equilibrate. The next day, media were aspirated, washed once with PBS, and replaced with different media as indicated. Proliferation was measured on IncuCyte S3 using phase object confluence as a readout.

### Metabolite sample preparation

Intracellular metabolite fractions were prepared from cells grown in 6-well plates. The media was aspirated, and cells were incubated with cold (−80°C) 80% methanol (1mL/well) on dry ice for 10 minutes. Then, the wells were scraped with a cell scraper and transferred to 1.5mL tubes on dry ice. To measure GlcNAc concentration in different conditioned media, 0.8mL of ice-cold 100% methanol was added to 0.2mL of conditioned media, and the mixture was incubated on dry ice for 10 minutes.

After incubation of cell or media fractions on dry ice, the tubes were centrifuged at 15,000rpm for 10 minutes at 4°C to pellet the insoluble material, and the supernatant was collected in a fresh 1.5mL tube. Metabolite levels of intercellular fractions were normalized to the protein content of a parallel sample, and all samples were lyophilized on a SpeedVac, and re-suspended in a 50:50 mixture of methanol and water in HPLC vials for LC-MS/MS analysis.

### Liquid chromatography-coupled mass spectrometry

To detect UDP-GlcNAc, the Shimadzu NEXERA integrated UHPLC system with a LC30AD pump, SIL30AC autosampler, CTO30A column oven, CBM20A controller was coupled with the AB Sciex TripleTOF 5600 MS system with DuoSpray ion source. All calibrations and operations were under control of Analyst TF 1.7.1. Calibrations of TOF-MS and TOF-MS/MS were achieved through reference APCI source of SCEIX calibration solution. A high throughput LC method of 8 min with flowrate of 0.4 ml/min with a Supelco Ascentis Express HILIC (75 mm X 3.0 mm, 2.7 µm). Solvent A was made of 20 mM ammonium acetate of 95% water and 5% acetonitrile at pH 9.0. Solvent B was 95% acetonitrile and 5% water. LC gradient 0.0-0.5 min 90% B, 3 min 50% B, 4.10 min 1% B, 5.5 min 1% B, 5.6 min 90% B, 6.5 min 90% B, 8 min stopping. Key parameters on the MS were the CE and CE spread of −35ev, 15ev. Data were compared to a reference standard. Data processing was performed by Sciex PeakView, MasterView, LibraryView and MQ software tools and ChemSpider database.

To measure GlcNAc concentration in the various conditioned media, we utilized an Agilent Technologies Triple Quad 6470 LC/MS system consisting of 1290 Infinity II LC Flexible Pump (Quaternary Pump), 1290 Infinity II Multisampler, 1290 Infinity II Multicolumn Thermostat with 6 port valve and 6470 triple quad mass spectrometer. Agilent Masshunter Workstation Software LC/MS Data Acquisition for 6400 Series Triple Quadrupole MS with Version B.08.02 is used for compound optimization and sample data acquisition.

A GlcNAc standard was used to establish parameters, against which conditioned media were analyzed. For LC, an Agilent ZORBAX RRHD Extend-C18, 2.1 × 150 mm, 1.8 um and ZORBAX Extend Fast Guards for UHPLC were used in the separation. LC gradient profile is: at 0.25 ml/min, 0-2.5 min, 100% A; 7.5 min, 80% A and 20% C; 13 min 55% A and 45% C; 20 min, 1% A and 99% C; 24 min, 1% A and 99% C; 24.05 min, 1% A and 99% D; 27 min, 1% A and 99% D; at 0.8 ml/min, 27.5-31.35 min, 1% A and 99% D; at 0.6 ml/min, 31.50 min, 1% A and 99% D; at 0.4 ml/min, 32.25-39.9 min, 100% A; at 0.25 ml/min, 40 min, 100% A. Column temp is kept at 35 °C, samples are at 4 °C, injection volume is 2 µl. Solvent A is 97% water and 3% methanol 15 mM acetic acid and 10 mM tributylamine at pH of 5. Solvent C is 15 mM acetic acid and 10 mM tributylamine in methanol. Washing Solvent D is acetonitrile. LC system seal washing solvent 90% water and 10% isopropanol, needle wash solvent 75% methanol, 25% water.

6470 Triple Quad MS is calibrated with ESI-L Low concentration Tuning mix. Source parameters: Gas temp 150 °C, Gas flow 10 l/min, Nebulizer 45 psi, Sheath gas temp 325 °C, Sheath gas flow 12 l/min, Capillary −2000 V, Delta EMV −200 V. Dynamic MRM scan type is used with 0.07 min peak width, acquisition time is 24 min. Delta retention time of plus and minus 1 min, fragmentor of 40 eV and cell accelerator of 5 eV are incorporated in the method.

### Xenograft studies

Animal experiments were conducted in accordance with the Office of Laboratory Animal Welfare and approved by the Institutional Animal Care and Use Committees of the University of Michigan. NOD-SCID gamma (NSG) mice (Jackson Laboratory), 6-10 weeks old of both sexes, were maintained in the facilities of the Unit for Laboratory Animal Medicine (ULAM) under specific pathogen-free conditions. Protocol#: PRO00008877.

Wild type TU8988T and two verified GFAT1 null clones (B9 and D10) were trypsinized and suspended at 1:1 ratio of DMEM (Gibco, 11965-092) cell suspension to Matrigel (Corning, 354234). 150-200μL were used per injection. Orthotopic tumors were established by injecting 0.5 x 10^6^ cells in 50μL of 1:1 DMEM to Matrigel mixture. The experiment lasted five weeks. For subcutaneous xenograft studies with the pooled populations or validated clones, tumors were established with 5 x 10^6^ cells in 200μL of 1:1 DMEM to Matrigel mixture.

Tumor size was measured with digital calipers two times per week. Tumor volume (V) was calculated as V = 1/2(length x width^2^). At endpoint, final tumor volume and mass were measured prior to processing. Tissue was snap-frozen in liquid nitrogen then stored at −80°C.

### Western blot analysis

After SDS-PAGE, proteins were transferred to PVDF membrane, blocked with 5% milk, and incubated with primary antibody overnight at 4°C. The membranes were washed with TBST, incubated with the appropriate horseradish peroxidase-conjugated secondary antibody for 1hr and visualized on Bio-Rad imager with enhanced chemiluminescence detection system or exposed on radiographic film.

### Antibodies

The following antibodies were used in this study: VINCULIN (Cell Signaling 13901), ACTIN (Santa Cruz sc-47778), GAPDH (Cell Signaling 5174), GFAT1 (Abcam 125069), NAGK (Atlas Antibodies HPA035207), O-GlcNAc (Abcam 2735), secondary anti-mouse-HRP (Cell Signaling 7076), and secondary anti-rabbit-HRP (Cell Signaling 7074).

### Detection and quantification of macropinocytosis

The macropinocytosis index was measured as previously described (59). In brief, cells were seeded on the coverslips in 24-well plate for 24 hours and serum-starved for 18 h. Cells were incubated with 1mg/ml high molecular weight TMR–dextran (Fina Biosolutions) in serum-free medium for 30 min at 37 °C. Cells were then washed 5 times with cold DPBS and fixed in 4% polyformaldehyde for 15 min. The coverslips were mounted onto slides using DAKO Mounting Medium (DAKO) in which nuclei were stained with DAPI. At least six images were captured for each sample using an Olympus FV1000 confocal microscope and analyzed using the particle analysis feature in ImageJ (NIH). The micropinocytosis index for each field was calculated as follow: Macropinocytosis Index = (total particle area/total cell area) × 100%.

### Hyaluronic acid, hyaluronidase, and heparin

Heparin was obtained from Sigma (H3393). Oligo HA (5kDa) was obtained from Lifecore Biomedical. Two different LMW HA were used in this study: 78 kDa HA (Pure Health solutions) and 60kDa HA (Lifecore Biomedical). To make 10mM oligo- or LMW HA media, HA was added slowly into high glucose DMEM without pyruvate, stirred for two hours at room temperature, and filtered through 0.20μm polyethersulfone membrane. FBS was added to a final concentration of 10%.

Hyaluronidase (Sigma H3506) treatment was performed as follows: 10mM LMW HA media and control media (DMEM + 10% FBS) were incubated with hyaluronidase, according to manufacturer’s instructions, overnight in a 37°C water bath. The next day, media were boiled for 15 minutes to denature hyaluronidase. The resulting media were mixed 1:1 with fresh growth media to avoid effects of nutrient/metabolite exhaustion.

### Preparation of necrotic FL5.**12 cells**

Necrotic FL5.12 cells were prepared as described previously (33). Cells were washed three times with PBS, cultured in the FL5.12 media without IL-3 (100 million cells/mL) for 72 hours. The necrotic cells were spun down at 13,000 rpm for 10 minutes at 4°C, and the pellets were stored at −80°C until use.

### Statistical analysis

Statistics were performed using GraphPad Prism 8. Groups of two were analyzed with two-tailed students t test. Groups of more than two were analyzed with one-way ANOVA Tukey post-hoc test. All error bars represent mean with standard deviation. A *P* value of less than 0.05 was considered statistically significant. All group numbers and explanation of significant values are presented within the figure legends.

## Acknowledgments

This work was funded by T32AI007413 and F31CA243344 (PK); K99CA241357 and P30DK034933 (CJH); T32AI007413 and F31CA24745701 (SAK); R01CA237466, R01CA252037 and R21CA212958 (KRK), StandUp2Cancer (KRK), Thompson Family Foundation (KRK), Geoffrey Beene Cancer Research Center at MSKCC and the STARR Cancer Consortium as well as the MSKCC NIH/NCI Cancer Center Support Core Grant P30CA008748; a Pancreatic Cancer Action Network/AACR Pathway to Leadership award (13-70-25-LYSS), Junior Scholar Award from The V Foundation for Cancer Research (V2016-009), Kimmel Scholar Award from the Sidney Kimmel Foundation for Cancer Research (SKF-16-005), a 2017 AACR NextGen Grant for Transformative Cancer Research (17-20-01-LYSS), the Cancer Center support grant (P30 CA046592); and NIH grants R37CA237421, R01CA248160, R01CA244931 (CAL). Metabolomics studies were supported by NIH grant DK097153, the Charles Woodson Research Fund, and the UM Pediatric Brain Tumor Initiative.

## SUPPLEMENTARY FIGURES

**Supplementary Figure 1.**
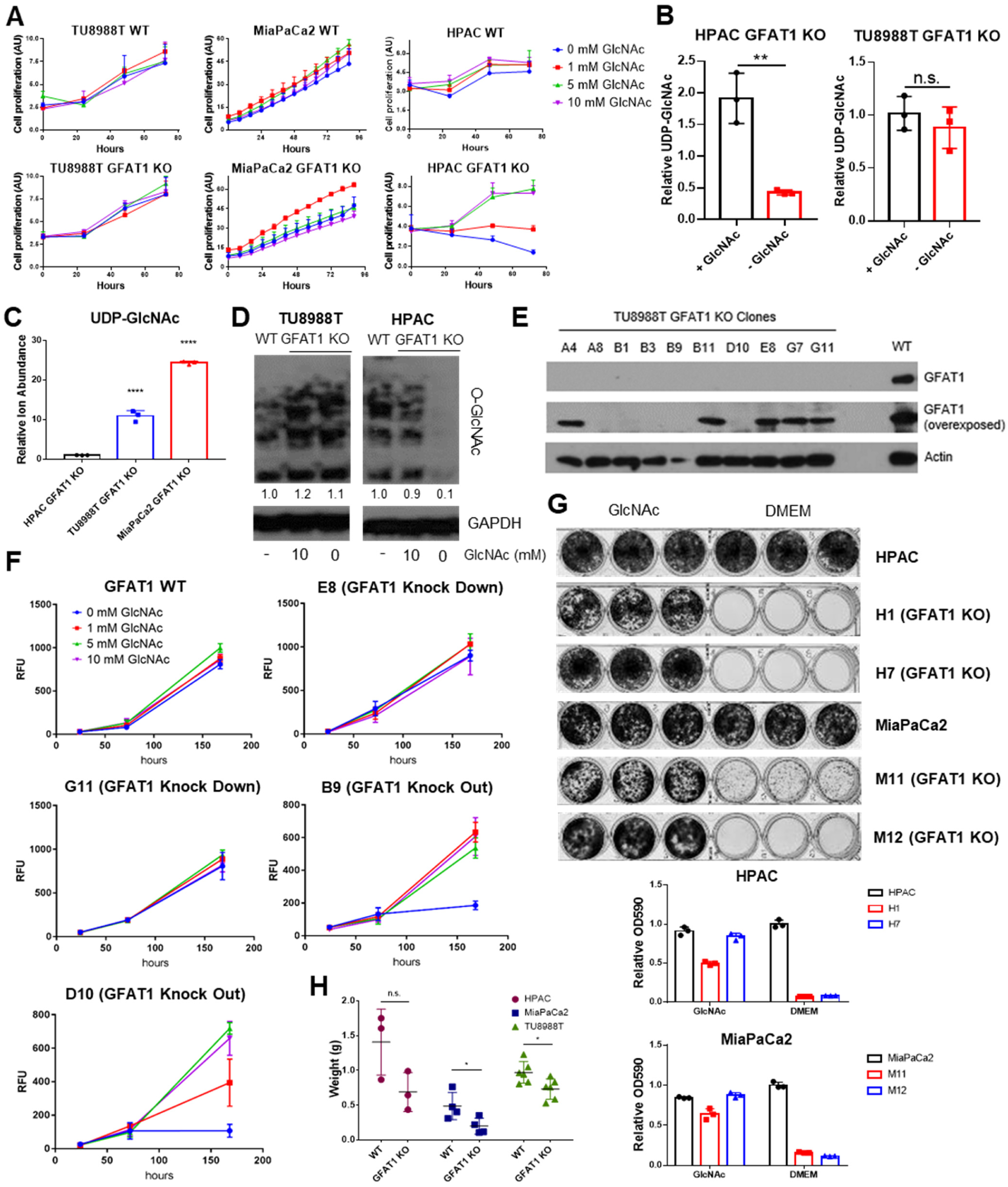
Additional characterization of GFAT1 knockout PDA populations and clonal lines. (**A**) Proliferation kinetics of parental PDA cell lines and corresponding pooled populations of GFAT1 knockout cells supplemented with varying concentrations of GlcNAc (n=3). Cell quantity was assessed by Cyquant (DNA intercalating dye) and plotted in absorbance units (AU). (**B,C**) UDP-GlcNAc levels measured by liquid chromatography-coupled tandem mass spectrometry (LC-MS/MS) in (**B**) TU8988T and HPAC GFAT1 knockout lines in the presence or absence of 10 mM GlcNAc for 3 days, presented as relative abundance (n=3), and (**C**) TU8988T, HPAC, and MiaPaCa2 GFAT1 knockout cells grown without GlcNAc for 3 days (n=3), presented as relative ion abundance. (**D**) Western blot of proteome O-GlcNAc and loading control GAPDH in parental and GFAT1 knockout TU8988T and HPAC. GFAT1 knockout lines were grown in the presence or absence of 10mM GlcNAc for 3 days. (**E**) Western blot for GFAT1, at short and long exposure, and ACTIN loading control in a panel of clonal cell lines selected from the pooled population of GFAT1 knockout TU8988T cells. (**F**) Proliferation kinetics of parental TU8988T (GFAT1 WT) and clonal cell lines E8, G11, B9, and D10 selected from the pooled GFAT1 knockout population supplemented with varying concentrations of GlcNAc (n=3). Clones correspond to those in the western blot in **E**. Cell quantity was assessed by cell titer glo and plotted in relative fluorescent units (RFU). (**G**) Representative wells from proliferation assay in parental and two GFAT1 knockout clonal HPAC and MiaPaCa2 cell lines. At bottom, data are quantitated by crystal violet extraction and measurement of optical density (OD) at 590nm, n=3. (**H**) Tumors from parental (n=3) and GFAT1 knockout (n=3) HPAC; parental (n=4) and GFAT1 knockout (n=4) MiaPaCa2; and parental (n=6) and GFAT1 knockout (n=6) TU8988T cell lines grown subcutaneously in immunocompromised mice. Error bars represent mean ± SD. n.s., non-significant; \**P* < 0.05; ** *P* <0.01; **** *P* <0.0001.

**Supplementary Figure 2.**
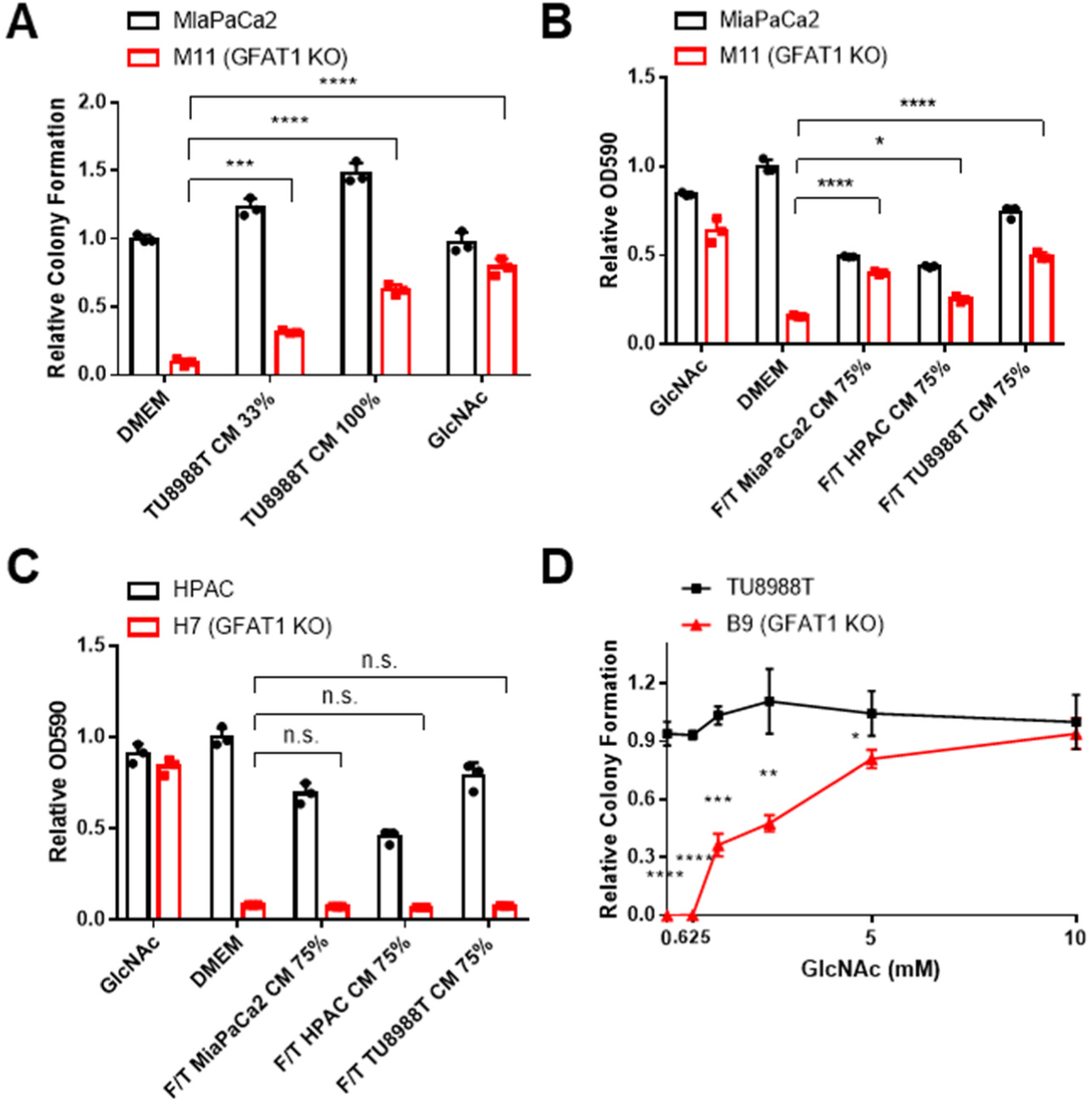
Characterization of rescue activity of conditioned media and GlcNAc. (**A**) Quantitation of colony forming assay data of parental MiaPaCa2 and GFAT1 knockout clonal line M11 in base media (DMEM), positive control GlcNAc, wild type TU8988T CM diluted 1:2 (33%) or used directly (100%) (n=3). (**B,C**) Quantitation of proliferation assay data of (**B**) parental MiaPaCa2 and GFAT1 knockout clonal line M11 and (**C**) parental HPAC and GFAT1 knockout clonal line H7 in base media (DMEM), positive control GlcNAc, or wild type TU8988T, HPAC, or MiaPaCa2 CM diluted 3:1 (75%) that was subjected to freeze-thaw (F/T) (n=3). Data represent crystal violet extracted from cells at endpoint and measured by optical density (OD) at 590nm. (**D**) GlcNAc dose response curve presented as relative colony number for parental TU8988T cells and GFAT1 knockout clonal line B9 (n=3). Error bars represent mean ± SD. n.s., non-significant; \**P* < 0.05; ** *P* <0.01; *** *P* <0.001; **** *P* <0.0001.

**Supplementary Figure 3.**
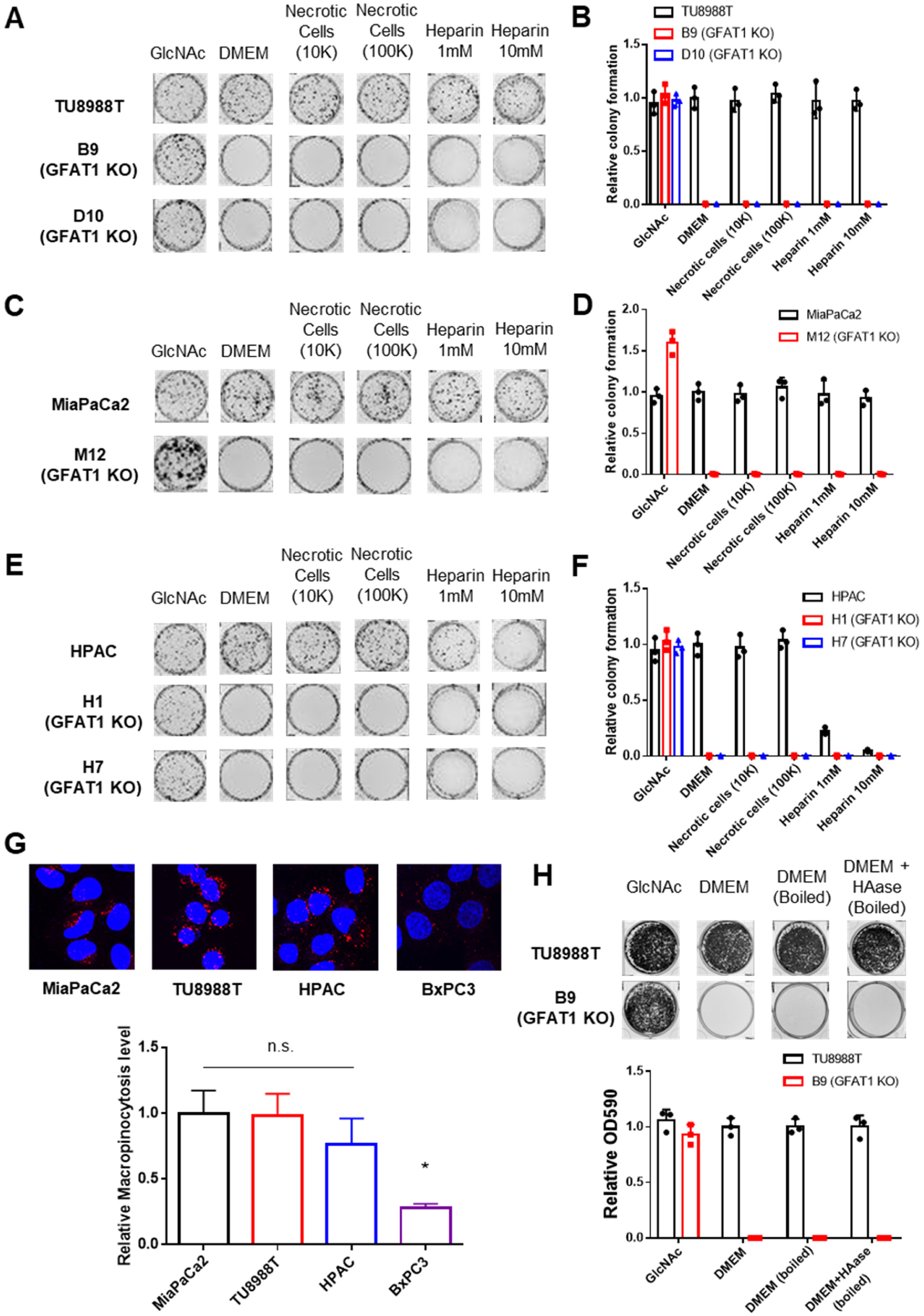
Characterization of macropinocytosis and glycosaminoglycan rescue activity in PDA and GFAT1 knockout cells. (**A-F**) Representative colony formation assays and their quantitation following treatment with two concentrations of heparin or necrotic cell debris that contain complete cellular contents, relative to base media (DMEM) and positive control GlcNAc in (**A,B**) parental and GFAT1 knockout TU8988T, (**C,D**) parental and GFAT1 knockout MiaPaCa2, and (**E,F**) parental and GFAT1 knockout HPAC. (**G**) Immunostaining images of intracellular fluorescently-tagged dextran (red) engulfed by macropinocytosis in PDA cell lines. Nuclear DAPI staining in blue. Quantitation of macropinocytotic index presented at bottom for n=6 wells per biological replicate (n=3). (**H**) Representative wells from a proliferation assay in parental TU8988T and GFAT1 knockout clone B9 in 10mM GlcNAc, base media (DMEM), base media supplemented 1:1 with boiled DMEM, or base media supplemented 1:1 with boiled HAase-treated DMEM. Data are quantitated below and represent crystal violet extracted from cells at endpoint and measured by optical density (OD) at 590nm (n=3). Error bars represent mean ± SD. n.s., non-significant; \**P* < 0.05.

